## Supplemental Materials for "General Trends in the Calnexin-Dependent Expression and Pharmacological Rescue of Clinical CFTR Variants"

**Contents**

-Figure S1
-Figure S2
-Figure S3
-Figure S4
-Figure S5
-Figure S6
-Figure S7
-Figure S8
-Figure S9
-Figure S10
-Table S1

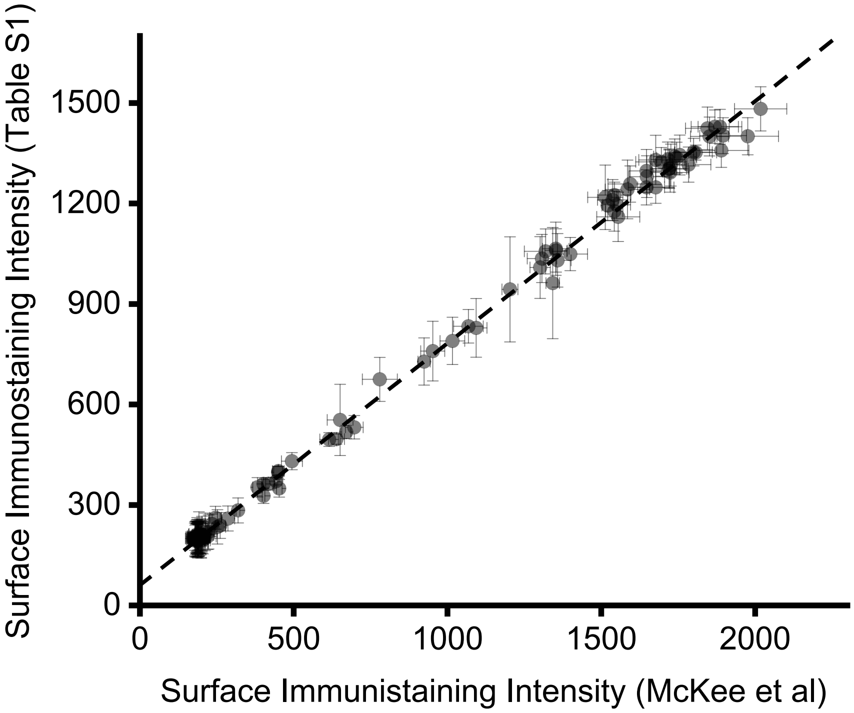


**Figure S1. Comparison of Deep Mutational Scanning Measurements Derived from First and Second Generation CF Variant Libraries.** A subset of surface immunostaining intensities for 129 CF variants derived from deep mutational scans using the library of 235 variants described herein are plotted against the corresponding 129 values that were previously reported from a deep mutational scan of the first generation library described in reference 5. Values represent the average of three biological replicates and error bars represent the standard deviation. A linear best fit is shown for reference (Pearson's R2 = 0.9975).

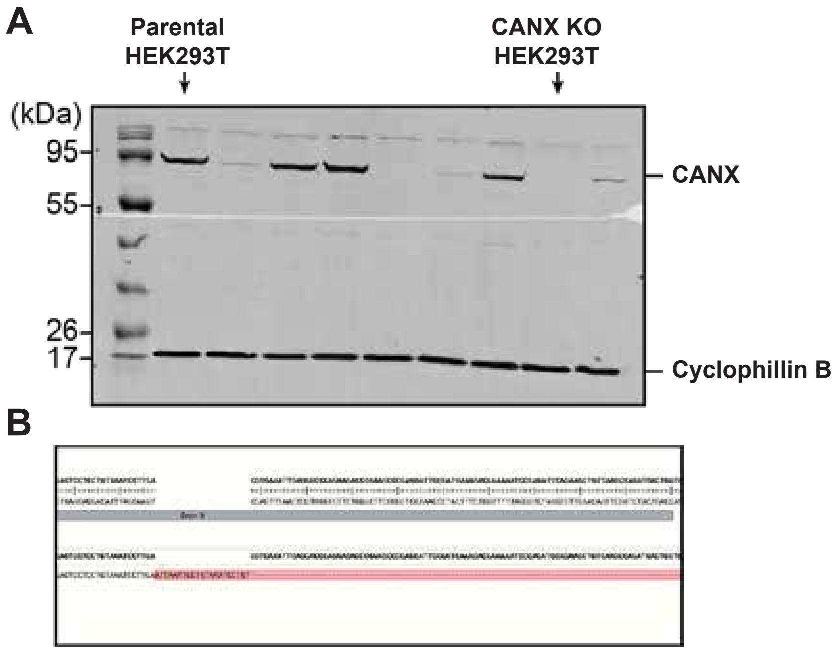


**Figure S2. Validation of Cas9 Mediated CANX Knockout Cells.** A) Western blot compares the levels of CANX protein expression across a series of clones isolated following transfection with a Cas9 RNP loaded with anti-*CANX* guide RNA. The lanes containing lystate from the parental cell line and the knockout line used for the studies detailed herein are indicated. An anti-cyclophillin B stain loading control is shown for reference. B) The aligment of a sanger sequencing read to the CANX gene sequence demonstrated that the chosen knockout clone indicated in panel A) contains a deletion within exon 8.

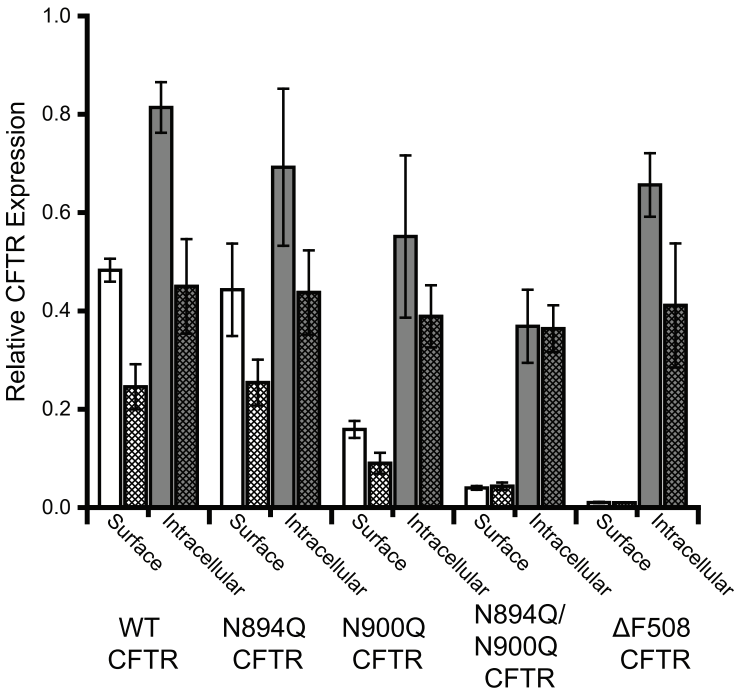


**Figure S3. Surface and Internal Expression of N-Glycan Knockouts in Parental and CANX KO Backgrounds.** A series of HA-tagged CFTR variants wer transiently expressed in parental HEK293T (open) and CANX KO (patterned) HEK 293T cells. Surface CFTR on intact cells was first immunostained with a Dylight-550 conjugated anti-HA antibody, then fixed and permeabilized prior to immunostaining of the intracellular CFTR with an Alexaflour-647 conjugated anti-HA antibody. Relative surface and intracellular immunostaining intensities were then measured by flow cytometry. A bar graph depicts the relative surface immunostaining intensity (white) and relative intracellular immunostaining intensity (gray) CFTR immunostaining for each CFTR variant. Values represent the average of three biological replicates and error bars represent the standard deviations. These data show that both WT and ΔF508 CFTR exhibit impaired expression in the absence of CANX. This expression defect persists for single mutants of either glycosylation site (N894 or N900). However, the CANX dependence of expression is lost for the double mutant that lacks N-linked glycosylation sites. Consistent with the findings in reference 19, these trends validate that CFTR exhibits lower expression in CANX knockout cells. Our results also suggest this expression defect depends upon its recognition of both N-linked glycans.

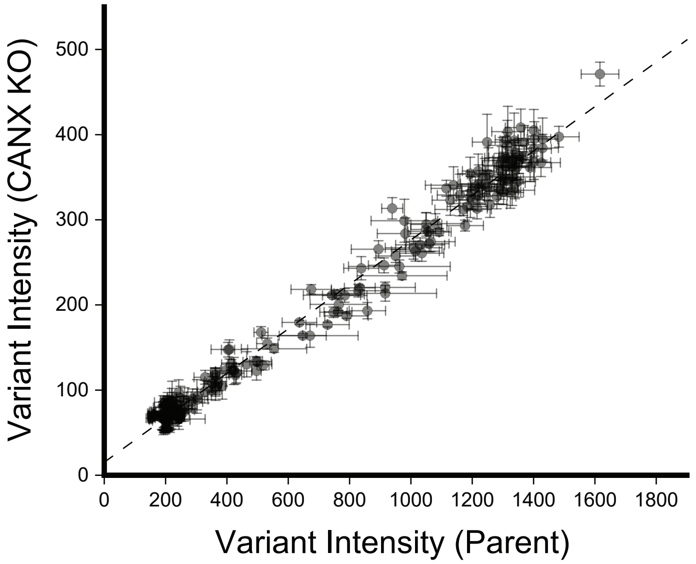


**Figure S4. Plasma Membrane Expression of CF Variants in Parental HEK293T Cells and CANX Knockout HEK293T Cells.** Deep mutational scanning measurements of the surface immunostaining intensities of 234 CF variants in parental HEK293T cells are plotted against the corresponding surface immunostaining intensities in CANX knockout cells. Values represent the average of three biological replicates and error bars represent the standard deviations. A line of best fit is shown for reference ($m$ = 0.26).

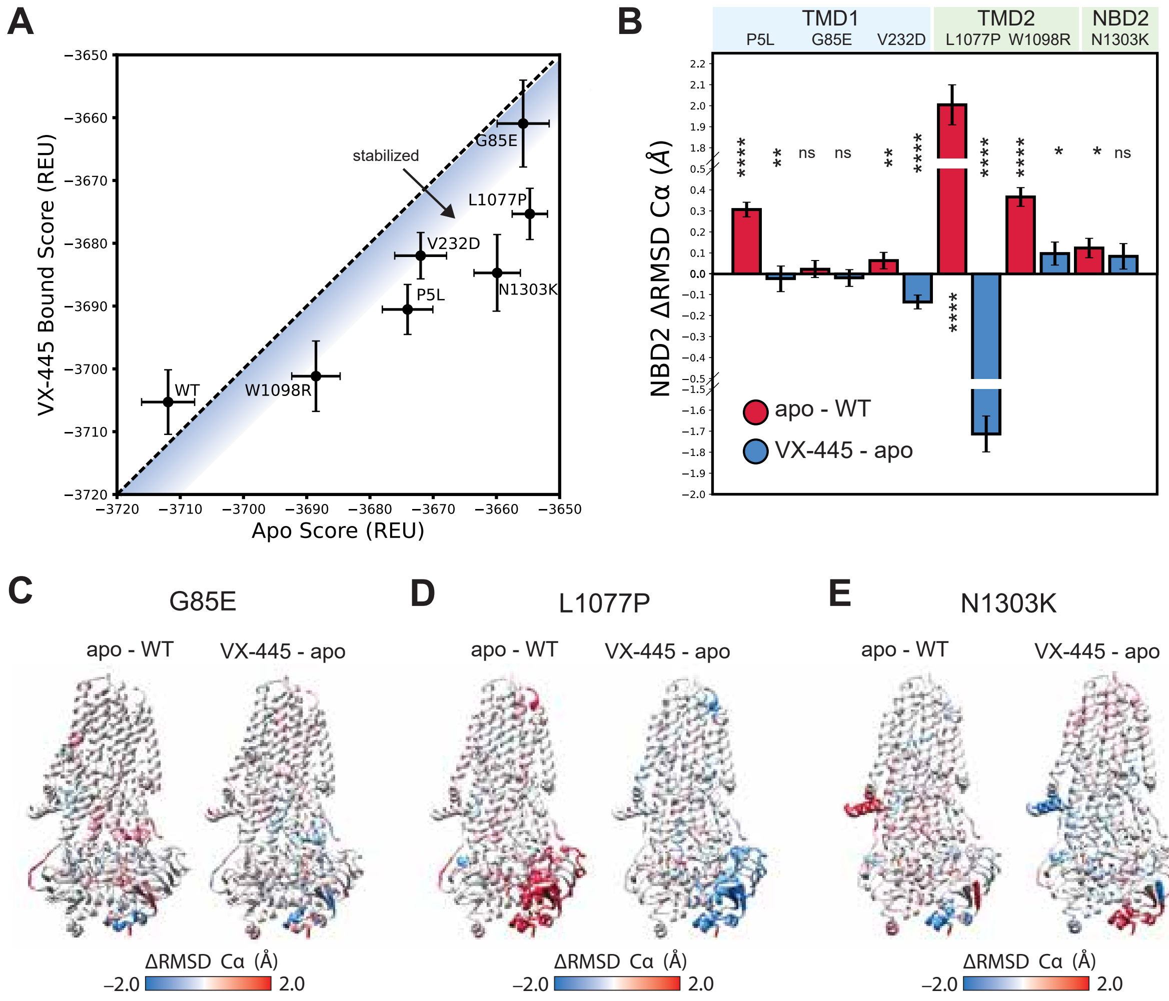


**Figure S5. VX-445-Mediated Suppression of Conformational Defects in NBD2.** Structural modeling was used to compare the conformational states of the apo and VX-445-bound active structures of WT CFTR and six rare CF variants (P5L, G85E, V232D, L1077P, W1098R, N1303K). A) The average Rosetta energy scores (± SEM) for the 100 lowest scoring models of the VX-445-bound state are plotted against those of the apo CFTR variant models. A reference line corresponding to no stabilization is shown for reference. All variants except for the non-responsive G85E fall below the line, which confirms VX-445 enhances stability. B) The total ΔRMSD of the active conformation of NBD2 is shown for variants bound to VX-445. Red bars show increasing deviations from the native NBD2 conformation in the mutant models and blue bars how much VX-445 suppresses these conformational defects in NBD2. C) Maps of the change in RMSD between G85E modeled with and without VX-445 shows which structural regions are stabilized by VX-445. Structurally variable regions in the ensemble shown in red, while areas adopting a more ordered conformation are shown in blue. VX-445 appears to primarily suppress conformational defects within the NBD2 region of this variant. D) Maps of the change in RMSD between L1077P modeled with and without VX-445 shows which structural regions are stabilized by VX-445. Structurally variable regions in the ensemble shown in red, while areas adopting a more ordered conformation are shown in blue. VX-445 appears to primarily suppress conformational defects within the NBD2 region of this variant. E) Maps of the change in RMSD between N1303K modeled with and without VX-445 shows that few structural regions are stabilized by VX-445 for N1303K, which responds poorly to VX-445 *in vitro*. Statistical significances were calculated using a non-parametric Wilcoxon signed-rank test compared to zero to determine if distributions changes were significantly different from zero, and p values were depicted by *< 0.05, **< 0.01, ***< 0.001, and ****< 0.0001. Error bars indicate the standard error of the mean.

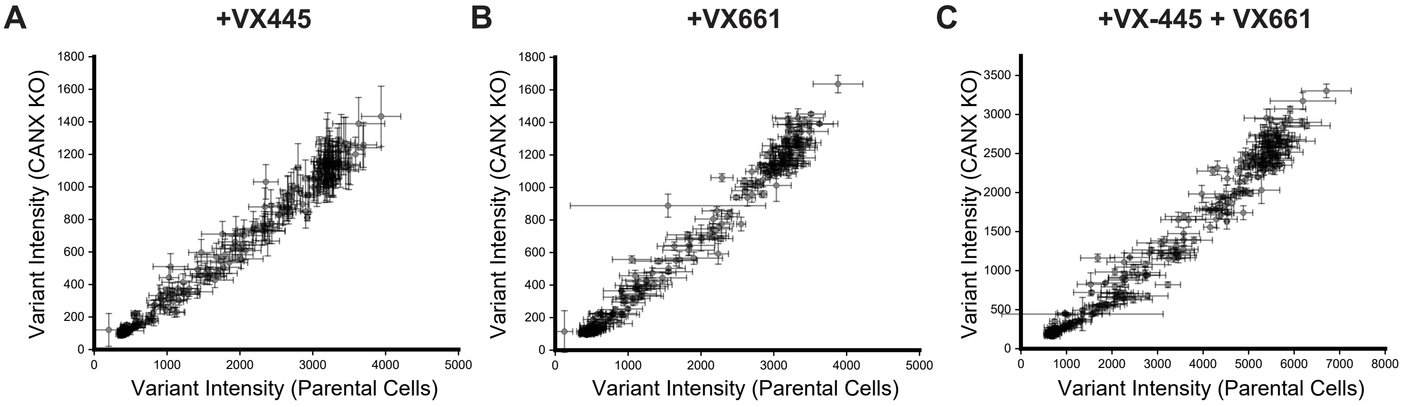


**Figure S6. Influence of Correctors on the Plasma Membrane Expression in Parental and CANX Knock-out Cells.** Deep mutational scanning measurements of the plasma membrane expression of CF variants in CANX knockout cells in the presence of A) 3 μM VX-445, B) 3 μM VX-661, or C) 3 μM VX-445 + 3 μM VX-661 are plotted against the corresponding variant measurements under identical conditions in the parental HEK-293T cell line. Values represent the average of three biological replicates and error bars represent the standard deviation. Trends are generally linear and there are few variants that stray from the tend, which suggests CF variants that respond in one cell line have a similar response in the other.

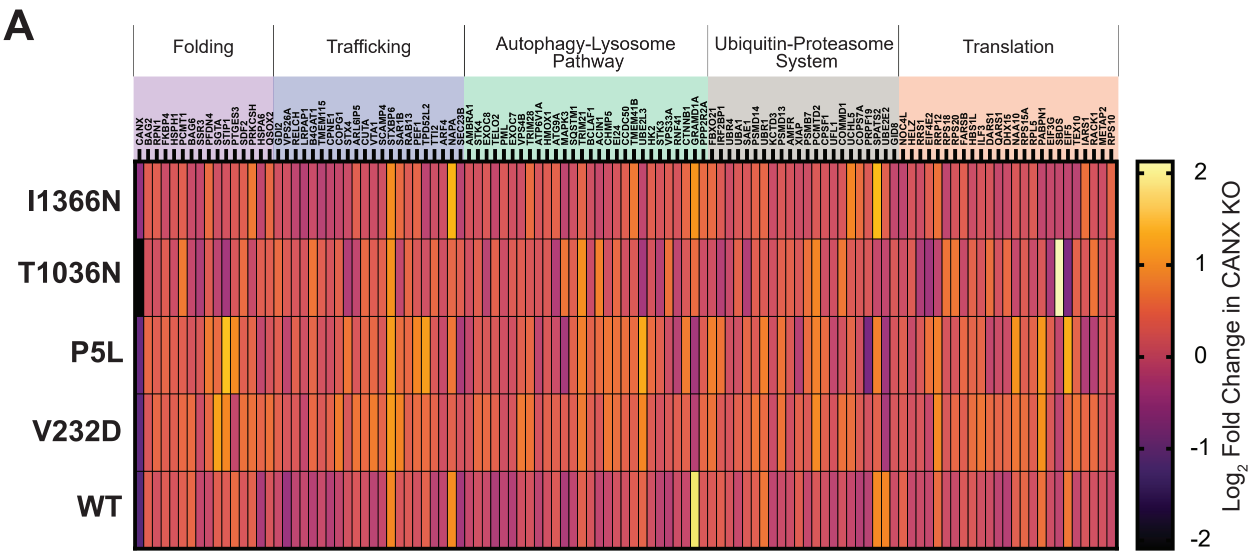


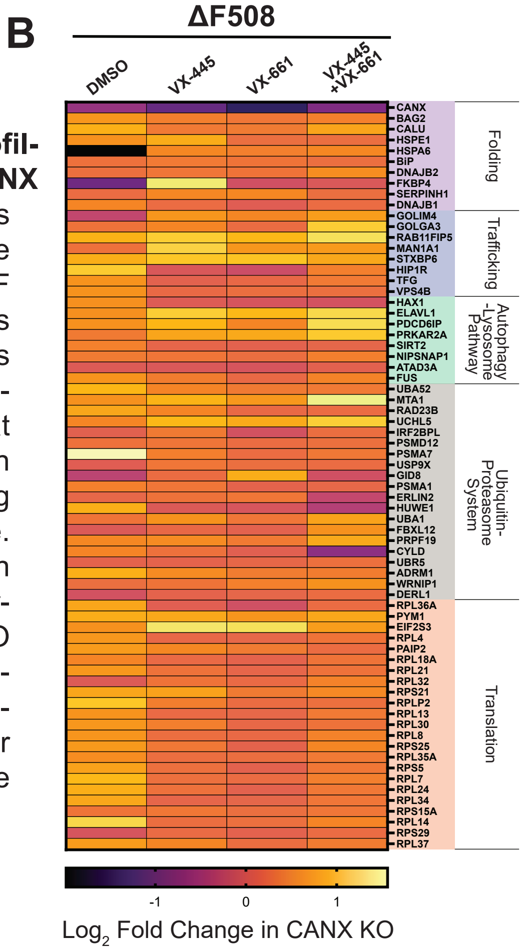


**Figure S7. Comparative Interactome Profiling of CF variants in Parental and CANX Knockout HEK293T Cells.** Proteomic mass spectrometry was used to compare the protein-protein interactions formed by CF variants in the context of CANX knockout cells relative to the parental cell line. A) Heat maps depict the $\log_2$ fold change in protein abundances of various classes of interactors that associate with each indicated CF variant in CANX KO cells relative to their corresponding abundance in the parental HEK293T cell line. B) Heat maps depict the $\log_2$ fold change in protein abundances of various classes of interactors that associate with ΔF508 in CANX KO cells treated with 3 μM of the indicated corrector(s) relative to their corresponding abundance in the parental HEK293T cell line under identical conditions. Protein interactors that are grouped according to their pathways.

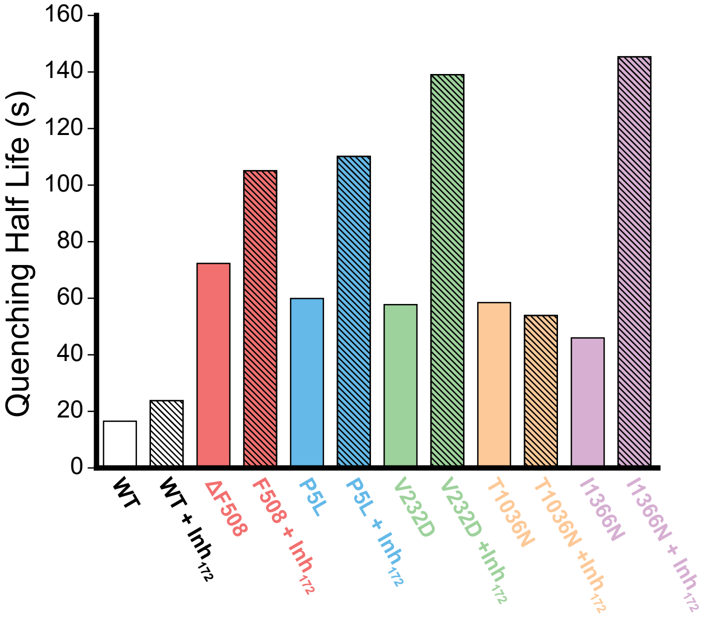


**Figure S8. Impact of a CFTR-Specific Inhibitor on Observed hYFP Quenching Kinetics in Recombinant Cells Expressing CF Variants.** The functional properties of CF variants are compared in various cells that feature endogenous CANX expression or deficient CANX expression under various experimental conditions. Bar graphs depict the fitted half-lives of the hYFP quenching reactions among cells expressing each indicated CF variant in parental HEK-293T cells treated with either vehicle (open bars) or with 10 μM of the CFTR-specific inhibitor-172 ($Inh_{172}$). A slowing of quenching was observed in the context of recombinant cell lines expressing each variant except T1036N. Though it is unclear why inhibition was not observed in this cell line, it is possible that this variant may have limited affinity for the inhibitor. Nevertheless, this cell line exhibits an intermediate quenching rate that lies between wild-type and ΔF508, which suggests the observed quencing is unlikely to be non-specific.

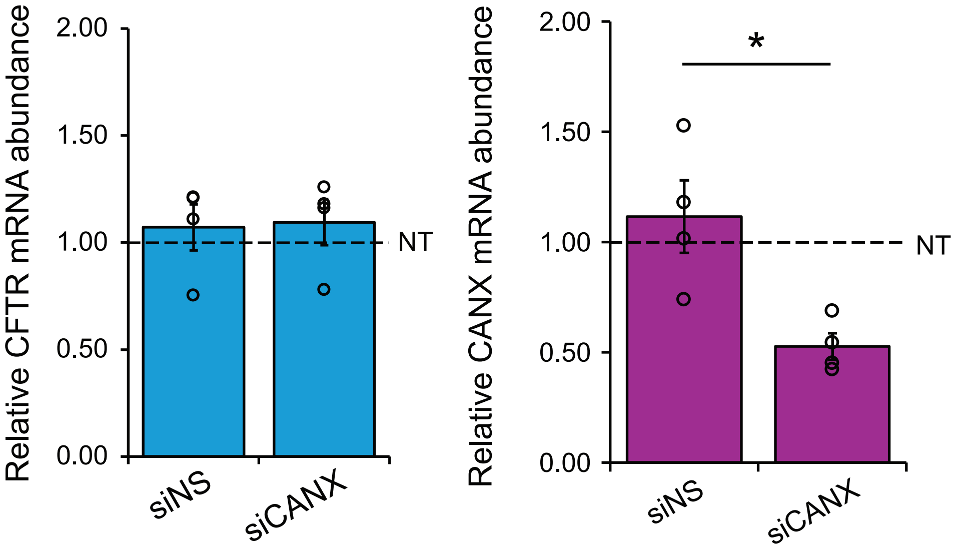


**Figure S9. Validation of siRNA-Mediated CANX Knockdowns in FRT Cells .** Quantitative reverse-transcriptase PCR was used to measure the relative abundance of the CFTR and CANX transcripts in FRT cells following transfection with either a non-specific or CANX-specific siRNA.ΔCT values for the target transcripts were normalized relative to the corresponding values of actin B transcript.

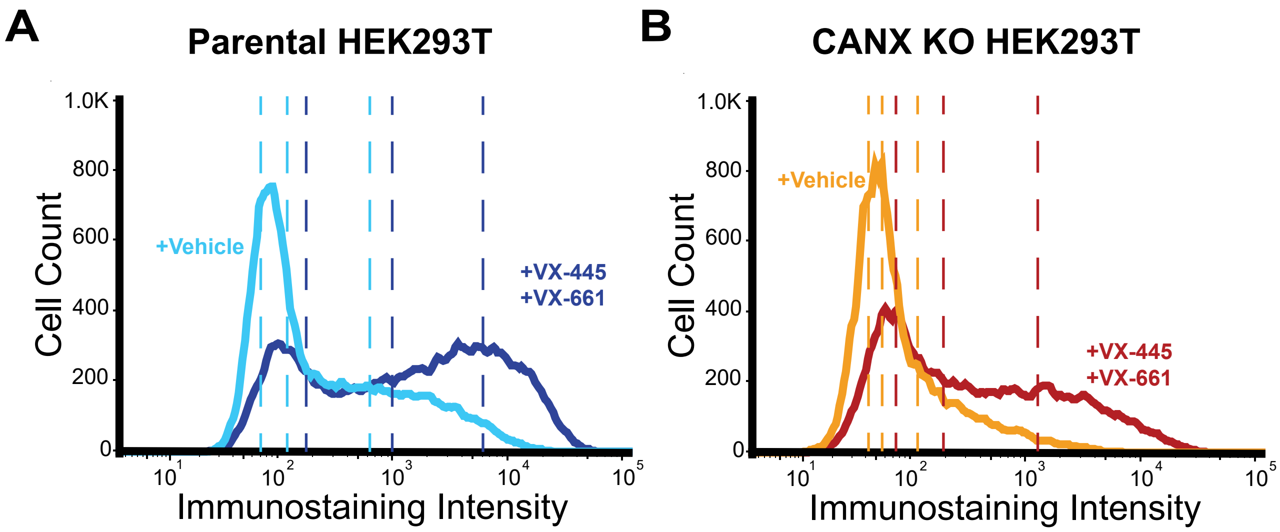


**Figure S10. Comparison of Sorting Gates for Deep Mutational Scanning.** Histograms depict the distribution of surface immunostaining intensities of recombinant libraries expressing CF variants in the A) parental and B) CANX knockout HEK293T cell lines. The approximate positions of the sorting gates used to fractionate each cell line into four equal quartiles based on immunostaining intensity are shown. Raw data and exact gating information for each experiment can be found in the Mendeley data directory.

### Table S1. Deep Mutational Scanning Measurements for CF Variants in Treated Parental and CANX KO Cells

| Variant | Amino Acid Substitution | CFTR sub-domain | Parental Surface Immuno-staining Intensity (Apo) | Parental Surface Immuno-staining Intensity (VX-661) | Parental Surface Immuno-staining Intensity (VX-445) | Parental Surface Immuno-staining Intensity (VX-445 + VX-661) | CANX KO Surface Immuno-staining Intensity (Apo) | CANX KO Surface Immuno-staining Intensity (VX-661) | CANX KO Surface Immuno-staining Intensity (VX-445) | CANX KO Surface Immuno-staining Intensity (VX-445 + VX-661) |
|---|---|---|---|---|---|---|---|---|---|---|
| **R3W** | 7A>T | Lasso | 913 ± 51 | 2602 ± 179 | 2934 ± 24 | 5281 ± 254 | 247 ± 9 | 980 ± 25 | 949 ± 119 | 2336 ± 48 |
| **P5L** | 14C>T | Lasso | 260 ± 37 | 921 ± 235 | 2438 ± 152 | 4780 ± 95 | 78 ± 15 | 219 ± 19 | 774 ± 123 | 2015 ± 95 |
| **S13F** | 38C>T | Lasso | 202 ± 43 | 498 ± 103 | 616 ± 51 | 2179 ± 291 | 72 ± 17 | 126 ± 13 | 169 ± 18 | 722 ± 73 |
| **L15P** | 44T>C | Lasso | 212 ± 49 | 566 ± 126 | 535 ± 12 | 1845 ± 336 | 72 ± 19 | 126 ± 9 | 134 ± 11 | 538 ± 54 |
| **G27R** | 79G>A | Lasso | 205 ± 43 | 518 ± 112 | 644 ± 52 | 2107 ± 395 | 70 ± 18 | 114 ± 21 | 154 ± 18 | 663 ± 24 |
| **R31C** | 91C>T | Lasso | 532 ± 35 | 1853 ± 338 | 2224 ± 259 | 4364 ± 369 | 155 ± 6 | 680 ± 43 | 737 ± 66 | 1934 ± 50 |
| **R31L** | 92G>T | Lasso | 352 ± 18 | 1219 ± 289 | 2343 ± 279 | 4274 ± 352 | 108 ± 11 | 378 ± 26 | 731 ± 86 | 1784 ± 11 |
| **A46D** | 137C>A | Lasso | 218 ± 44 | 564 ± 128 | 1461 ± 213 | 3573 ± 348 | 70 ± 15 | 133 ± 11 | 455 ± 50 | 1474 ± 73 |
| **E56K** | 166G>A | Lasso | 208 ± 48 | 825 ± 188 | 446 ± 42 | 1723 ± 325 | 72 ± 16 | 249 ± 16 | 115 ± 17 | 557 ± 23 |
| **W57G** | 169T>G | Lasso | 200 ± 48 | 450 ± 85 | 392 ± 39 | 637 ± 109 | 67 ± 17 | 103 ± 15 | 102 ± 22 | 180 ± 40 |
| **E60K** | 178G>A | Lasso | 198 ± 45 | 513 ± 117 | 388 ± 37 | 903 ± 161 | 66 ± 14 | 130 ± 18 | 100 ± 20 | 263 ± 16 |
| **P67L** | 200C>T | Lasso | 212 ± 50 | 786 ± 146 | 447 ± 48 | 1782 ± 293 | 70 ± 15 | 225 ± 19 | 110 ± 16 | 557 ± 57 |
| **R74W** | 220C>T | Lasso | 518 ± 20 | 1703 ± 233 | 1562 ± 220 | 3568 ± 322 | 129 ± 5 | 556 ± 49 | 443 ± 13 | 1274 ± 36 |
| **R74Q** | 221G>A | Lasso | 784 ± 73 | 2975 ± 174 | 2175 ± 172 | 5170 ± 152 | 211 ± 8 | 1142 ± 35 | 722 ± 65 | 2387 ± 40 |
| **R75Q** | 224G>A | Lasso | 759 ± 89 | 2886 ± 76 | 1938 ± 232 | 5045 ± 53 | 212 ± 6 | 1137 ± 9 | 662 ± 81 | 2330 ± 105 |
| **G85E** | 254G>A | TMD1 | 200 ± 43 | 454 ± 103 | 384 ± 42 | 676 ± 97 | 67 ± 15 | 109 ± 16 | 102 ± 17 | 184 ± 25 |
| **G91R** | 271G>A | TMD1 | 195 ± 47 | 528 ± 131 | 409 ± 38 | 1639 ± 312 | 67 ± 17 | 121 ± 13 | 105 ± 16 | 456 ± 35 |
| **E92K** | 274G>A | TMD1 | 204 ± 57 | 488 ± 89 | 405 ± 49 | 720 ± 93 | 70 ± 14 | 112 ± 15 | 105 ± 15 | 190 ± 27 |
| **Q98R** | 293A>G | TMD1 | 210 ± 47 | 951 ± 173 | 490 ± 21 | 2170 ± 408 | 69 ± 15 | 296 ± 23 | 124 ± 13 | 708 ± 94 |
| **P99L** | 296C>T | TMD1 | 428 ± 20 | 906 ± 212 | 1415 ± 211 | 1921 ± 357 | 118 ± 10 | 226 ± 8 | 355 ± 40 | 562 ± 30 |
| **L102R** | 305T>G | TMD1 | 199 ± 42 | 441 ± 96 | 381 ± 43 | 697 ± 161 | 64 ± 15 | 96 ± 15 | 90 ± 14 | 159 ± 28 |
| **Y109N** | 325T>A | TMD1 | 743 ± 93 | 2484 ± 77 | 1939 ± 188 | 4444 ± 160 | 211 ± 2 | 939 ± 13 | 641 ± 91 | 1941 ± 15 |
| **D110Y** | 328G>T | TMD1 | 495 ± 20 | 2140 ± 204 | 1594 ± 190 | 4397 ± 231 | 134 ± 5 | 721 ± 25 | 464 ± 44 | 1715 ± 55 |
| **D110H** | 328G>C | TMD1 | 1065 ± 79 | 3097 ± 102 | 2606 ± 163 | 5337 ± 143 | 274 ± 12 | 1111 ± 36 | 845 ± 75 | 2285 ± 59 |
| **D110E** | 330C>A | TMD1 | 1337 ± 46 | 3511 ± 195 | 3237 ± 85 | 5912 ± 341 | 393 ± 32 | 1452 ± 10 | 1197 ± 146 | 3071 ± 34 |
| **E116K** | 346G>A | TMD1 | 982 ± 92 | 2990 ± 37 | 2343 ± 223 | 4936 ± 68 | 284 ± 18 | 1195 ± 49 | 877 ± 116 | 2491 ± 79 |
| **R117C** | 349C>T | TMD1 | 833 ± 50 | 2405 ± 122 | 2310 ± 148 | 4593 ± 102 | 220 ± 2 | 841 ± 23 | 731 ± 90 | 1887 ± 45 |
| **R117H** | 350G>A | TMD1 | 1060 ± 65 | 2853 ± 51 | 2668 ± 56 | 4953 ± 141 | 272 ± 10 | 959 ± 21 | 839 ± 67 | 2010 ± 46 |
| **R117P** | 350G>C | TMD1 | 979 ± 109 | 2679 ± 132 | 2656 ± 93 | 4776 ± 74 | 299 ± 25 | 1023 ± 21 | 952 ± 118 | 2280 ± 51 |
| **R117L** | 350G>T | TMD1 | 762 ± 45 | 2191 ± 188 | 2080 ± 210 | 4156 ± 144 | 192 ± 1 | 701 ± 25 | 609 ± 57 | 1557 ± 63 |
| **A120T** | 358G>A | TMD1 | 1302 ± 43 | 3206 ± 142 | 3217 ± 57 | 5466 ± 178 | 346 ± 15 | 1212 ± 46 | 1046 ± 108 | 2453 ± 74 |
| **G126D** | 377G>A | TMD1 | 350 ± 26 | 1472 ± 330 | 1303 ± 157 | 3413 ± 340 | 99 ± 11 | 444 ± 33 | 374 ± 47 | 1313 ± 42 |
| **I148T** | 443T>C | TMD1 | 1049 ± 49 | 3134 ± 194 | 2588 ± 30 | 5261 ± 210 | 289 ± 20 | 1165 ± 29 | 874 ± 109 | 2401 ± 36 |
| **Y161D** | 481T>G | TMD1 | 199 ± 51 | 474 ± 108 | 427 ± 50 | 1665 ± 232 | 69 ± 16 | 120 ± 9 | 110 ± 18 | 535 ± 34 |
| **Y161C** | 482A>C | TMD1 | 206 ± 43 | 582 ± 121 | 749 ± 35 | 3240 ± 362 | 72 ± 17 | 142 ± 10 | 204 ± 23 | 1152 ± 78 |
| **L165S** | 494T>C | TMD1 | 199 ± 58 | 495 ± 104 | 806 ± 17 | 2573 ± 266 | 73 ± 15 | 126 ± 19 | 224 ± 34 | 955 ± 120 |
| **R170H** | 509G>A | TMD1 | 1216 ± 33 | 3030 ± 104 | 2953 ± 56 | 5204 ± 129 | 313 ± 12 | 1096 ± 10 | 952 ± 107 | 2198 ± 21 |
| **L172I** | 514C>A | TMD1 | 1116 ± 50 | 2953 ± 114 | 3109 ± 130 | 5344 ± 384 | 337 ± 9 | 1219 ± 79 | 1155 ± 138 | 2651 ± 129 |
| **G178R** | 532G>A | TMD1 | 1009 ± 92 | 2784 ± 73 | 2655 ± 101 | 4991 ± 172 | 267 ± 13 | 1025 ± 26 | 875 ± 85 | 2224 ± 21 |
| **G178E** | 533G>A | TMD1 | 1369 ± 32 | 3348 ± 165 | 3392 ± 128 | 5817 ± 362 | 391 ± 20 | 1369 ± 23 | 1277 ± 167 | 2892 ± 95 |
| **F191V** | 571T>G | TMD1 | 233 ± 42 | 787 ± 168 | 1526 ± 206 | 3575 ± 369 | 77 ± 16 | 220 ± 9 | 466 ± 65 | 1397 ± 73 |
| **D192G** | 575A>G | TMD1 | 251 ± 39 | 753 ± 194 | 1774 ± 241 | 3356 ± 364 | 78 ± 15 | 172 ± 12 | 506 ± 53 | 1160 ± 51 |
| **E193K** | 577G>A | TMD1 | 838 ± 57 | 2607 ± 136 | 2426 ± 94 | 4887 ± 101 | 243 ± 14 | 1003 ± 18 | 885 ± 97 | 2346 ± 50 |

| Variant | Amino Acid Substitution | CFTR sub-domain | Parental Surface Immuno-staining Intensity (Apo) | Parental Surface Immuno-staining Intensity (VX-661) | Parental Surface Immuno-staining Intensity (VX-445) | Parental Surface Immuno-staining Intensity (VX-445 + VX-661) | CANX KO Surface Immuno-staining Intensity (Apo) | CANX KO Surface Immuno-staining Intensity (VX-661) | CANX KO Surface Immuno-staining Intensity (VX-445) | CANX KO Surface Immuno-staining Intensity (VX-445 + VX-661) |
|---|---|---|---|---|---|---|---|---|---|---|
| **G194R** | 580G>A | TMD1 | 765 ± 93 | 2159 ± 269 | 2537 ± 190 | 4477 ± 196 | 201 ± 7 | 746 ± 33 | 832 ± 99 | 1832 ± 100 |
| **G194V** | 581G>T | TMD1 | 303 ± 36 | 1255 ± 286 | 1233 ± 194 | 3342 ± 464 | 89 ± 12 | 422 ± 54 | 352 ± 52 | 1257 ± 36 |
| **H199Y** | 595C>T | TMD1 | 201 ± 45 | 454 ± 109 | 391 ± 47 | 689 ± 122 | 70 ± 16 | 111 ± 12 | 104 ± 20 | 188 ± 27 |
| **V201M** | 601G>A | TMD1 | 496 ± 15 | 2282 ± 166 | 1701 ± 262 | 4523 ± 37 | 123 ± 11 | 776 ± 88 | 468 ± 34 | 1631 ± 99 |
| **P205S** | 613C>T | TMD1 | 205 ± 41 | 478 ± 97 | 437 ± 22 | 841 ± 154 | 71 ± 14 | 123 ± 15 | 112 ± 23 | 245 ± 16 |
| **L206W** | 617T>G | TMD1 | 203 ± 46 | 618 ± 81 | 556 ± 19 | 1586 ± 203 | 67 ± 14 | 151 ± 19 | 138 ± 17 | 454 ± 18 |
| **L227R** | 680T>G | TMD1 | 200 ± 48 | 476 ± 121 | 401 ± 72 | 657 ± 157 | 70 ± 14 | 118 ± 11 | 105 ± 18 | 197 ± 32 |
| **V232D** | 695T>A | TMD1 | 202 ± 47 | 513 ± 113 | 446 ± 35 | 1117 ± 209 | 69 ± 13 | 120 ± 10 | 108 ± 21 | 272 ± 11 |
| **A234D** | 701C>A | TMD1 | 423 ± 24 | 1266 ± 281 | 1549 ± 310 | 3080 ± 417 | 124 ± 7 | 390 ± 21 | 495 ± 92 | 1219 ± 31 |
| **Q237E** | 709C>G | TMD1 | 1299 ± 68 | 3076 ± 72 | 3282 ± 109 | 5328 ± 189 | 347 ± 17 | 1129 ± 22 | 1127 ± 119 | 2377 ± 47 |
| **Q237H** | 711G>C | TMD1 | 1339 ± 34 | 3401 ± 162 | 3487 ± 166 | 5954 ± 306 | 343 ± 19 | 1229 ± 36 | 1131 ± 112 | 2678 ± 90 |
| **R258G** | 772A>G | TMD1 | 239 ± 56 | 663 ± 138 | 749 ± 41 | 1551 ± 227 | 77 ± 15 | 148 ± 22 | 173 ± 20 | 440 ± 33 |
| **M265R** | 794T>G | TMD1 | 223 ± 41 | 568 ± 185 | 571 ± 33 | 1069 ± 146 | 75 ± 15 | 130 ± 19 | 146 ± 17 | 325 ± 7 |
| **F311L** | 933C>G | TMD1 | 1199 ± 42 | 2912 ± 94 | 3141 ± 26 | 5260 ± 110 | 324 ± 12 | 1094 ± 13 | 1047 ± 93 | 2352 ± 21 |
| | 933C>A | TMD1 | 1194 ± 40 | 2901 ± 89 | 3169 ± 22 | 5428 ± 106 | 354 ± 19 | 1197 ± 20 | 1165 ± 140 | 2637 ± 116 |
| **G314E** | 941G>A | TMD1 | 894 ± 89 | 2598 ± 98 | 2703 ± 150 | 5091 ± 116 | 265 ± 10 | 1041 ± 17 | 975 ± 115 | 2489 ± 63 |
| **L320V** | 958T>G | TMD1 | 1401 ± 21 | 3491 ± 325 | 3425 ± 186 | 5637 ± 494 | 405 ± 24 | 1389 ± 26 | 1251 ± 176 | 2711 ± 80 |
| **R334W** | 1000C>T | TMD1 | 1429 ± 50 | 3421 ± 211 | 3339 ± 136 | 5746 ± 483 | 395 ± 25 | 1407 ± 28 | 1214 ± 148 | 2838 ± 65 |
| **R334L** | 1001G>T | TMD1 | 219 ± 51 | 555 ± 111 | 602 ± 25 | 1298 ± 236 | 76 ± 15 | 143 ± 13 | 148 ± 13 | 342 ± 13 |
| **R334Q** | 1001G>A | TMD1 | 1310 ± 76 | 2985 ± 91 | 3277 ± 87 | 5342 ± 243 | 363 ± 23 | 1127 ± 10 | 1153 ± 144 | 2454 ± 53 |
| **I336K** | 1007T>A | TMD1 | 203 ± 44 | 537 ± 109 | 469 ± 33 | 1194 ± 229 | 73 ± 18 | 130 ± 17 | 125 ± 14 | 351 ± 19 |
| **T338I** | 1013C>T | TMD1 | 1194 ± 45 | 3206 ± 193 | 3098 ± 100 | 5322 ± 316 | 327 ± 24 | 1177 ± 21 | 1080 ± 116 | 2466 ± 83 |
| **S341P** | 1021T>C | TMD1 | 363 ± 18 | 1354 ± 250 | 1290 ± 164 | 3454 ± 370 | 100 ± 13 | 402 ± 29 | 340 ± 29 | 1167 ± 66 |
| **L346P** | 1037T>C | TMD1 | 188 ± 52 | 463 ± 84 | 387 ± 44 | 1144 ± 184 | 67 ± 13 | 114 ± 11 | 99 ± 13 | 321 ± 9 |
| **R347H** | 1040G>A | TMD1 | 1401 ± 55 | 3350 ± 219 | 3697 ± 180 | 5851 ± 609 | 368 ± 20 | 1182 ± 19 | 1258 ± 138 | 2658 ± 17 |
| **R347P** | 1040G>C | TMD1 | 239 ± 39 | 877 ± 182 | 1104 ± 141 | 2784 ± 445 | 72 ± 15 | 163 ± 20 | 228 ± 29 | 676 ± 34 |
| | 1040G>T | TMD1 | 858 ± 59 | 2237 ± 140 | 2925 ± 28 | 4885 ± 206 | 193 ± 10 | 592 ± 63 | 811 ± 20 | 1742 ± 89 |
| **A349V** | 1046C>T | TMD1 | 1036 ± 72 | 3029 ± 63 | 2664 ± 100 | 5321 ± 122 | 261 ± 10 | 1082 ± 28 | 866 ± 60 | 2274 ± 55 |
| **R352W** | 1054C>T | TMD1 | 511 ± 23 | 1636 ± 235 | 1470 ± 174 | 3457 ± 385 | 168 ± 7 | 639 ± 26 | 597 ± 80 | 1653 ± 95 |
| **R352Q** | 1055G>A | TMD1 | 1424 ± 63 | 3292 ± 82 | 3515 ± 134 | 5674 ± 277 | 367 ± 16 | 1132 ± 37 | 1138 ± 125 | 2350 ± 15 |
| **Q359K/ T360K** | 1075-1079C>A | TMD1 | 327 ± 23 | 1184 ± 267 | 1583 ± 235 | 3463 ± 393 | 98 ± 13 | 337 ± 25 | 450 ± 51 | 1197 ± 44 |
| **Q359R** | 1076A>G | TMD1 | 1184 ± 66 | 2943 ± 121 | 3011 ± 58 | 5257 ± 169 | 316 ± 18 | 1102 ± 39 | 998 ± 93 | 2317 ± 40 |
| **W361R** | 1081T>C | TMD1 | 194 ± 49 | 435 ± 87 | 427 ± 64 | 738 ± 80 | 69 ± 15 | 106 ± 16 | 110 ± 13 | 203 ± 29 |
| **S364P** | 1090T>C | TMD1 | 209 ± 43 | 457 ± 103 | 423 ± 42 | 708 ± 113 | 67 ± 15 | 108 ± 13 | 101 ± 15 | 184 ± 25 |
| **G404R** | 1210G>C | | 1312 ± 42 | 3342 ± 232 | 3212 ± 84 | 5476 ± 342 | 372 ± 19 | 1268 ± 8 | 1131 ± 134 | 2576 ± 75 |
| **D443Y** | 1327G>T | NBD1 | 362 ± 15 | 1162 ± 276 | 1071 ± 154 | 2736 ± 419 | 121 ± 5 | 378 ± 14 | 342 ± 34 | 1079 ± 40 |
| **L453S** | 1358T>C | NBD1 | 244 ± 85 | 527 ± 43 | 499 ± 182 | 948 ± 153 | 66 ± 12 | 112 ± 12 | 115 ± 23 | 255 ± 25 |
| **A455E** | 1364C>A | NBD1 | 197 ± 50 | 482 ± 140 | 400 ± 63 | 726 ± 95 | 67 ± 17 | 101 ± 10 | 97 ± 15 | 185 ± 37 |
| **V456F** | 1366G>T | NBD1 | 203 ± 44 | 426 ± 95 | 408 ± 30 | 777 ± 116 | 67 ± 15 | 103 ± 12 | 101 ± 15 | 215 ± 24 |
| **V456A** | 1367T>C | NBD1 | 198 ± 51 | 460 ± 91 | 398 ± 46 | 756 ± 91 | 71 ± 19 | 120 ± 12 | 105 ± 16 | 210 ± 43 |
| **G463D** | 1388G>T | NBD1 | 200 ± 33 | 515 ± 119 | 731 ± 74 | 2288 ± 392 | 69 ± 14 | 119 ± 18 | 180 ± 22 | 705 ± 81 |
| **L467P** | 1400T>C | NBD1 | 189 ± 48 | 410 ± 95 | 367 ± 50 | 640 ± 87 | 67 ± 14 | 104 ± 10 | 95 ± 16 | 180 ± 34 |
| **M470V** | 1408A>G | NBD1 | 1315 ± 63 | 2987 ± 143 | 2798 ± 36 | 4316 ± 210 | 387 ± 29 | 1256 ± 35 | 1119 ± 147 | 2317 ± 88 |
| **E474K** | 1420G>A | NBD1 | 196 ± 44 | 434 ± 92 | 452 ± 21 | 1269 ± 283 | 67 ± 13 | 105 ± 14 | 109 ± 14 | 363 ± 6 |
| **G480S** | 1438G>T | NBD1 | 206 ± 41 | 450 ± 101 | 438 ± 21 | 1050 ± 168 | 71 ± 18 | 113 ± 14 | 118 ± 18 | 296 ± 29 |
| **S492F** | 1475C>T | NBD1 | 195 ± 50 | 428 ± 91 | 374 ± 35 | 734 ± 105 | 67 ± 16 | 107 ± 15 | 100 ± 14 | 196 ± 19 |
| **I502T** | 1505T>C | NBD1 | 199 ± 43 | 426 ± 81 | 375 ± 57 | 700 ± 156 | 68 ± 16 | 109 ± 15 | 105 ± 16 | 208 ± 31 |

| Variant | Amino Acid Substitution | CFTR sub-domain | Parental Surface Immuno-staining Intensity (Apo) | Parental Surface Immuno-staining Intensity (VX-661) | Parental Surface Immuno-staining Intensity (VX-445) | Parental Surface Immuno-staining Intensity (VX-445 + VX-661) | CANX KO Surface Immuno-staining Intensity (Apo) | CANX KO Surface Immuno-staining Intensity (VX-661) | CANX KO Surface Immuno-staining Intensity (VX-445) | CANX KO Surface Immuno-staining Intensity (VX-445 + VX-661) |
|---|---|---|---|---|---|---|---|---|---|---|
| **F508C** | 1523T>G | NBD1 | 1015 ± 78 | 2970 ± 108 | 3056 ± 46 | 5590 ± 283 | 263 ± 12 | 1090 ± 30 | 964 ± 85 | 2480 ± 53 |
| **D513G** | 1538A>G | NBD1 | 199 ± 46 | 449 ± 101 | 404 ± 39 | 736 ± 124 | 69 ± 15 | 108 ± 14 | 103 ± 20 | 213 ± 30 |
| **V520F** | 1558G>T | NBD1 | 201 ± 49 | 445 ± 98 | 380 ± 48 | 681 ± 131 | 67 ± 16 | 107 ± 13 | 97 ± 14 | 192 ± 30 |
| **E528E** | 1584G>A | NBD1 | 1349 ± 57 | 3303 ± 171 | 3281 ± 85 | 5641 ± 349 | 345 ± 18 | 1148 ± 32 | 1054 ± 106 | 2414 ± 33 |
| **S549N** | 1646G>A | NBD1 | 1340 ± 53 | 3197 ± 138 | 3227 ± 62 | 5604 ± 178 | 379 ± 14 | 1311 ± 40 | 1179 ± 134 | 2792 ± 86 |
| **S549R** | 1645A>C | NBD1 | 404 ± 21 | 1097 ± 215 | 1027 ± 133 | 2265 ± 376 | 147 ± 9 | 458 ± 33 | 442 ± 65 | 1110 ± 117 |
| | 1647T>A | NBD1 | 415 ± 9 | 1108 ± 204 | 1073 ± 120 | 2264 ± 339 | 131 ± 5 | 393 ± 20 | 366 ± 42 | 914 ± 55 |
| | 1647T>G | NBD1 | 385 ± 10 | 1068 ± 165 | 1036 ± 80 | 2251 ± 567 | 105 ± 15 | 291 ± 17 | 256 ± 44 | 661 ± 88 |
| **G551S** | 1651G>A | NBD1 | 1483 ± 66 | 3627 ± 252 | 3690 ± 250 | 6283 ± 509 | 398 ± 12 | 1391 ± 9 | 1241 ± 135 | 2855 ± 34 |
| **G551D** | 1652G>A | NBD1 | 1429 ± 52 | 3489 ± 257 | 3588 ± 129 | 6172 ± 428 | 384 ± 16 | 1344 ± 47 | 1201 ± 104 | 2900 ± 48 |
| **R553N** | 1658G>A | NBD1 | 1334 ± 31 | 3197 ± 156 | 3179 ± 148 | 5442 ± 153 | 371 ± 22 | 1251 ± 31 | 1112 ± 119 | 2594 ± 72 |
| **A554E** | (1661C>A) | NBD1 | 329 ± 37 | 968 ± 175 | 918 ± 73 | 2063 ± 298 | 115 ± 8 | 313 ± 26 | 313 ± 35 | 792 ± 34 |
| **L558S** | 1673T>C | NBD1 | 193 ± 47 | 443 ± 72 | 365 ± 41 | 639 ± 114 | 67 ± 14 | 103 ± 14 | 99 ± 16 | 170 ± 29 |
| **A559T** | 1675G>A | NBD1 | 199 ± 51 | 440 ± 82 | 398 ± 46 | 659 ± 81 | 66 ± 17 | 102 ± 17 | 95 ± 20 | 160 ± 21 |
| **R560K** | 1679G>A | NBD1 | 196 ± 51 | 425 ± 96 | 368 ± 39 | 614 ± 109 | 66 ± 15 | 97 ± 13 | 91 ± 19 | 170 ± 25 |
| **R560T** | 1679G>C | NBD1 | 202 ± 49 | 468 ± 111 | 396 ± 27 | 669 ± 91 | 66 ± 14 | 104 ± 14 | 97 ± 18 | 187 ± 33 |
| **R560S** | 1680A>C | NBD1 | 192 ± 49 | 448 ± 119 | 382 ± 46 | 666 ± 94 | 70 ± 17 | 108 ± 10 | 103 ± 19 | 189 ± 32 |
| **A561E** | 1682C>A | NBD1 | 194 ± 51 | 410 ± 71 | 376 ± 45 | 659 ± 79 | 69 ± 17 | 117 ± 17 | 107 ± 13 | 193 ± 25 |
| **V562I** | 1684G>A | NBD1 | 1057 ± 67 | 2698 ± 128 | 2646 ± 52 | 4692 ± 233 | 286 ± 1 | 958 ± 48 | 877 ± 85 | 2016 ± 35 |
| **Y563N** | 1687T>A | NBD1 | 193 ± 50 | 423 ± 85 | 364 ± 44 | 656 ± 63 | 71 ± 23 | 107 ± 20 | 98 ± 29 | 196 ± 44 |
| **Y563D** | 1687T>G | NBD1 | 193 ± 50 | 452 ± 127 | 388 ± 40 | 654 ± 81 | 69 ± 13 | 107 ± 18 | 103 ± 12 | 192 ± 31 |
| **Y569D** | 1705T>G | NBD1 | 206 ± 51 | 457 ± 90 | 408 ± 42 | 682 ± 99 | 69 ± 17 | 114 ± 13 | 103 ± 19 | 197 ± 40 |
| **P574H** | 1721C>A | NBD1 | 203 ± 50 | 475 ± 109 | 440 ± 45 | 1047 ± 184 | 69 ± 17 | 110 ± 14 | 118 ± 19 | 303 ± 12 |
| **F575Y** | 1724T>A | NBD1 | 227 ± 49 | 571 ± 130 | 567 ± 23 | 1353 ± 213 | 81 ± 11 | 161 ± 13 | 171 ± 17 | 494 ± 20 |
| **G576A** | 1727G>C | NBD1 | 1400 ± 58 | 3318 ± 138 | 3352 ± 92 | 5659 ± 291 | 383 ± 15 | 1297 ± 55 | 1152 ± 103 | 2685 ± 30 |
| **D579G** | 1736A>G | NBD1 | 398 ± 18 | 1102 ± 213 | 1000 ± 98 | 2227 ± 304 | 123 ± 9 | 378 ± 6 | 351 ± 34 | 902 ± 84 |
| **E588V** | 1763A>T | NBD1 | 417 ± 8 | 1136 ± 253 | 1009 ± 123 | 2088 ± 333 | 124 ± 8 | 346 ± 39 | 316 ± 39 | 713 ± 44 |
| **S589T** | 1766G>C | NBD1 | 1170 ± 92 | 2920 ± 37 | 2832 ± 21 | 4928 ± 129 | 312 ± 7 | 1073 ± 23 | 963 ± 110 | 2163 ± 50 |
| **I601F** | 1801A>T | NBD1 | 236 ± 47 | 644 ± 156 | 550 ± 34 | 1089 ± 165 | 79 ± 17 | 142 ± 23 | 142 ± 16 | 330 ± 16 |
| **H609R** | 1826A>G | NBD1 | 201 ± 45 | 470 ± 93 | 421 ± 54 | 768 ± 123 | 72 ± 19 | 107 ± 19 | 100 ± 17 | 199 ± 24 |
| **A613T** | 1837G>A | NBD1 | 208 ± 46 | 476 ± 98 | 413 ± 43 | 773 ± 126 | 69 ± 13 | 110 ± 20 | 108 ± 14 | 209 ± 28 |
| **D614G** | 1841A>G | NBD1 | 211 ± 55 | 538 ± 123 | 448 ± 35 | 948 ± 139 | 74 ± 18 | 127 ± 12 | 126 ± 16 | 295 ± 9 |
| **I618T** | 1853T>C | NBD1 | 209 ± 46 | 488 ± 110 | 471 ± 52 | 994 ± 192 | 69 ± 14 | 110 ± 12 | 115 ± 14 | 270 ± 11 |
| **G622D** | 1865G>A | NBD1 | 232 ± 40 | 723 ± 187 | 906 ± 75 | 2722 ± 328 | 77 ± 15 | 185 ± 5 | 263 ± 44 | 974 ± 81 |
| **G628R** | 1882G>A | NBD1 | 203 ± 54 | 489 ± 143 | 419 ± 44 | 812 ± 145 | 69 ± 16 | 113 ± 13 | 110 ± 16 | 234 ± 30 |
| | 1882G>C | NBD1 | 196 ± 50 | 443 ± 79 | 398 ± 47 | 760 ± 138 | 73 ± 16 | 119 ± 18 | 122 ± 21 | 252 ± 32 |
| **R668C** | 2002C>T | R Domain | 1210 ± 61 | 2968 ± 149 | 2900 ± 39 | 5071 ± 152 | 339 ± 11 | 1144 ± 10 | 1051 ± 114 | 2482 ± 64 |
| **S737F** | 2210C>T | R Domain | 1365 ± 46 | 3295 ± 83 | 3273 ± 39 | 5655 ± 202 | 373 ± 7 | 1287 ± 33 | 1141 ± 108 | 2608 ± 18 |
| **P750L** | 2249C>T | R Domain | 363 ± 13 | 1192 ± 292 | 1228 ± 219 | 2757 ± 423 | 104 ± 11 | 332 ± 17 | 341 ± 36 | 940 ± 19 |
| **R751L** | 2252G>T | R Domain | 1199 ± 34 | 3102 ± 89 | 3010 ± 57 | 5095 ± 211 | 334 ± 21 | 1147 ± 35 | 1045 ± 110 | 2386 ± 59 |
| **V754M** | 2260G>A | R Domain | 1316 ± 53 | 3192 ± 165 | 3189 ± 147 | 5394 ± 490 | 403 ± 30 | 1421 ± 37 | 1260 ± 156 | 2954 ± 114 |
| **I807M** | 2374C>G | R Domain | 1329 ± 63 | 3229 ± 153 | 3251 ± 150 | 5529 ± 300 | 370 ± 21 | 1249 ± 26 | 1136 ± 115 | 2625 ± 55 |
| **E822K** | 2464G>A | R Domain | 636 ± 56 | 1830 ± 354 | 2184 ± 142 | 4064 ± 249 | 179 ± 2 | 614 ± 34 | 745 ± 104 | 1796 ± 79 |
| **D836Y** | 2506G>T | | 1050 ± 50 | 2749 ± 112 | 2715 ± 111 | 4808 ± 100 | 295 ± 14 | 1017 ± 22 | 915 ± 107 | 2139 ± 71 |
| **S912L** | 2735C>T | TMD2 | 939 ± 34 | 2289 ± 152 | 2356 ± 168 | 4212 ± 345 | 313 ± 13 | 1060 ± 24 | 1031 ± 106 | 2275 ± 54 |
| **D924N** | 2770G>A | TMD2 | 1312 ± 37 | 3331 ± 349 | 3630 ± 357 | 6192 ± 717 | 394 ± 8 | 1428 ± 55 | 1388 ± 161 | 3173 ± 109 |
| **L927P** | 2780T>C | TMD2 | 675 ± 66 | 1813 ± 277 | 1760 ± 287 | 3579 ± 408 | 218 ± 6 | 709 ± 19 | 710 ± 78 | 1694 ± 44 |
| **R933G** | 2797A>G | TMD2 | 1617 ± 61 | 3882 ± 341 | 3938 ± 269 | 6709 ± 547 | 471 ± 14 | 1636 ± 54 | 1433 ± 186 | 3302 ± 87 |

| Variant | Amino Acid Substitution | CFTR sub-domain | Parental Surface Immuno-staining Intensity (Apo) | Parental Surface Immuno-staining Intensity (VX-661) | Parental Surface Immuno-staining Intensity (VX-445) | Parental Surface Immuno-staining Intensity (VX-445 + VX-661) | CANX KO Surface Immuno-staining Intensity (Apo) | CANX KO Surface Immuno-staining Intensity (VX-661) | CANX KO Surface Immuno-staining Intensity (VX-445) | CANX KO Surface Immuno-staining Intensity (VX-445 + VX-661) |
|---|---|---|---|---|---|---|---|---|---|---|
| **H939R** | 2816A>G | TMD2 | 1285 ± 63 | 3413 ± 85 | 3494 ± 135 | 5857 ± 264 | 337 ± 18 | 1145 ± 18 | 1106 ± 123 | 2573 ± 39 |
| **S945L** | 2834C>T | TMD2 | 284 ± 37 | 997 ± 201 | 1163 ± 112 | 2738 ± 306 | 86 ± 12 | 253 ± 10 | 303 ± 29 | 941 ± 68 |
| **M952T** | 2855T>C | TMD2 | 1241 ± 51 | 3020 ± 48 | 3115 ± 39 | 5421 ± 252 | 352 ± 20 | 1229 ± 10 | 1126 ± 142 | 2580 ± 72 |
| **M952I** | 2856G>A | TMD2 | 916 ± 99 | 2552 ± 61 | 2903 ± 68 | 5042 ± 182 | 221 ± 6 | 775 ± 42 | 849 ± 102 | 1989 ± 37 |
| **L967S** | 2900T>C | TMD2 | 728 ± 71 | 1902 ± 393 | 1852 ± 255 | 3415 ± 399 | 177 ± 3 | 566 ± 38 | 534 ± 67 | 1230 ± 46 |
| **G970R** | 2908G>C | TMD2 | 1358 ± 51 | 3228 ± 234 | 3225 ± 151 | 5314 ± 278 | 408 ± 22 | 1387 ± 25 | 1221 ± 141 | 2743 ± 126 |
| **G970D** | 2909G>A | TMD2 | 1160 ± 74 | 2821 ± 45 | 2738 ± 66 | 4828 ± 147 | 329 ± 24 | 1136 ± 16 | 1000 ± 110 | 2413 ± 105 |
| **S977F** | 2930C>T | TMD2 | 1218 ± 96 | 3010 ± 95 | 3276 ± 82 | 5622 ± 261 | 328 ± 18 | 1119 ± 47 | 1128 ± 124 | 2742 ± 130 |
| **I980K** | 2939T>A | TMD2 | 224 ± 39 | 605 ± 135 | 667 ± 34 | 1348 ± 216 | 73 ± 15 | 134 ± 12 | 150 ± 15 | 378 ± 3 |
| **L997F** | 2991G>C | TMD2 | 1297 ± 64 | 3275 ± 112 | 3451 ± 125 | 5910 ± 374 | 370 ± 16 | 1342 ± 28 | 1270 ± 158 | 2934 ± 83 |
| **Y1014C** | 3041A>G | TMD2 | 1139 ± 92 | 3111 ± 41 | 3083 ± 19 | 5309 ± 97 | 341 ± 22 | 1198 ± 37 | 1097 ± 124 | 2535 ± 93 |
| **F1016S** | 3047T>C | TMD2 | 361 ± 8 | 1334 ± 322 | 1677 ± 295 | 3682 ± 376 | 118 ± 9 | 475 ± 31 | 569 ± 85 | 1650 ± 93 |
| **I1027T** | 3080T>C | TMD2 | 1030 ± 80 | 2640 ± 108 | 2540 ± 50 | 4681 ± 132 | 270 ± 8 | 936 ± 51 | 824 ± 79 | 1970 ± 12 |
| **Y1032C** | 3095A>G | TMD2 | 234 ± 51 | 758 ± 242 | 1944 ± 177 | 3971 ± 288 | 82 ± 15 | 209 ± 10 | 712 ± 106 | 1982 ± 110 |
| **T1036N** | 3107C>A | TMD2 | 214 ± 66 | 587 ± 71 | 1135 ± 98 | 3228 ± 275 | 65 ± 15 | 124 ± 15 | 231 ± 29 | 820 ± 36 |
| **F1052V** | 3154T>G | TMD2 | 1248 ± 48 | 3204 ± 151 | 3205 ± 152 | 5471 ± 335 | 391 ± 33 | 1387 ± 51 | 1282 ± 173 | 2945 ± 125 |
| **T1053I** | 3158C>T | TMD2 | 1302 ± 56 | 3122 ± 105 | 3074 ± 72 | 5343 ± 109 | 333 ± 15 | 1163 ± 10 | 999 ± 100 | 2378 ± 50 |
| **H1054D** | 3160C>G | TMD2 | 207 ± 52 | 557 ± 107 | 1086 ± 139 | 3412 ± 304 | 69 ± 16 | 124 ± 16 | 293 ± 38 | 1215 ± 82 |
| **K1060T** | 3179A>C | TMD2 | 1129 ± 59 | 2870 ± 115 | 2783 ± 83 | 4958 ± 59 | 323 ± 18 | 1111 ± 10 | 992 ± 127 | 2331 ± 42 |
| **G1061R** | 3181G>C | TMD2 | 202 ± 42 | 469 ± 76 | 546 ± 57 | 1709 ± 321 | 69 ± 15 | 106 ± 16 | 139 ± 14 | 511 ± 28 |
| **R1066C** | 3196C>T | TMD2 | 194 ± 50 | 426 ± 68 | 363 ± 66 | 681 ± 106 | 67 ± 14 | 96 ± 12 | 94 ± 17 | 169 ± 30 |
| **R1066H** | 3197G>A | TMD2 | 207 ± 43 | 508 ± 112 | 834 ± 88 | 2822 ± 553 | 80 ± 18 | 152 ± 14 | 276 ± 29 | 1225 ± 126 |
| **A1067T** | 3199G>A | TMD2 | 747 ± 45 | 2361 ± 137 | 2405 ± 212 | 4870 ± 92 | 191 ± 6 | 820 ± 35 | 727 ± 75 | 2007 ± 50 |
| **G1069R** | 3205G>A | TMD2 | 1330 ± 73 | 3170 ± 138 | 3182 ± 19 | 5586 ± 205 | 335 ± 18 | 1135 ± 25 | 1013 ± 97 | 2344 ± 14 |
| **R1070W** | 3208C>T | TMD2 | 431 ± 25 | 1553 ± 331 | 1656 ± 253 | 3800 ± 405 | 120 ± 11 | 482 ± 11 | 427 ± 46 | 1390 ± 41 |
| **R1070Q** | 3209G>A | TMD2 | 1402 ± 54 | 3379 ± 169 | 3362 ± 133 | 5549 ± 317 | 395 ± 20 | 1344 ± 34 | 1221 ± 167 | 2734 ± 85 |
| **F1074L** | 3222T>G | TMD2 | 371 ± 11 | 1557 ± 229 | 1998 ± 205 | 4373 ± 180 | 107 ± 8 | 502 ± 32 | 618 ± 79 | 1860 ± 56 |
| **L1077P** | 3230T>C | TMD2 | 205 ± 46 | 458 ± 90 | 557 ± 17 | 2067 ± 314 | 68 ± 17 | 102 ± 16 | 144 ± 14 | 662 ± 59 |
| **H1085P** | 3254A>C | TMD2 | 202 ± 45 | 449 ± 84 | 423 ± 17 | 941 ± 123 | 69 ± 16 | 111 ± 18 | 110 ± 17 | 252 ± 20 |
| **H1085R** | 3254A>G | TMD2 | 214 ± 42 | 557 ± 139 | 803 ± 66 | 2258 ± 382 | 69 ± 17 | 123 ± 15 | 190 ± 22 | 639 ± 10 |
| **W1098R** | 3292T>C | TMD2 | 210 ± 47 | 481 ± 102 | 404 ± 46 | 676 ± 147 | 72 ± 17 | 129 ± 12 | 111 ± 18 | 207 ± 41 |
| **W1098C** | 3294G>C | TMD2 | 209 ± 45 | 550 ± 125 | 818 ± 64 | 2465 ± 422 | 68 ± 15 | 122 ± 12 | 175 ± 29 | 626 ± 39 |
| | 3294G>T | TMD2 | 221 ± 38 | 650 ± 204 | 818 ± 87 | 2293 ± 377 | 88 ± 23 | 189 ± 20 | 271 ± 34 | 880 ± 61 |
| **F1099L** | 3297C>A | TMD2 | 288 ± 31 | 1067 ± 234 | 2032 ± 263 | 4356 ± 177 | 90 ± 12 | 318 ± 4 | 643 ± 99 | 1771 ± 100 |
| **M1101K** | 3302T>A | TMD2 | 202 ± 47 | 463 ± 126 | 406 ± 45 | 740 ± 158 | 69 ± 15 | 122 ± 9 | 109 ± 20 | 211 ± 36 |
| **M1101R** | 3302T>G | TMD2 | 223 ± 30 | 509 ± 174 | 412 ± 51 | 745 ± 234 | 80 ± 14 | 152 ± 7 | 132 ± 21 | 271 ± 25 |
| **S1118F** | 3353C>T | TMD2 | 499 ± 46 | 1660 ± 291 | 2015 ± 303 | 4293 ± 457 | 133 ± 5 | 551 ± 34 | 557 ± 59 | 1658 ± 112 |
| **I1139V** | 3415A>G | TMD2 | 1222 ± 60 | 3124 ± 128 | 3023 ± 65 | 5157 ± 227 | 342 ± 22 | 1181 ± 39 | 1052 ± 134 | 2434 ± 175 |
| **D1152H** | 3454G>C | TMD2 | 1177 ± 59 | 3034 ± 205 | 2861 ± 77 | 5283 ± 400 | 293 ± 6 | 1012 ± 98 | 893 ± 50 | 2031 ± 171 |
| **V1153E** | 3458T>A | TMD2 | 302 ± 34 | 804 ± 170 | 980 ± 77 | 1858 ± 269 | 93 ± 20 | 203 ± 32 | 246 ± 26 | 566 ± 33 |
| **L1156F** | 3600G>T-3468G>T | TMD2 | 1248 ± 53 | 3224 ± 138 | 3263 ± 95 | 5550 ± 398 | 339 ± 16 | 1203 ± 2 | 1139 ± 142 | 2585 ± 71 |
| **S1159P** | 3475T>C | TMD2 | 1092 ± 39 | 2850 ± 96 | 3265 ± 131 | 5485 ± 328 | 286 ± 4 | 1046 ± 42 | 1076 ± 138 | 2493 ± 81 |
| **S1159F** | 3476C>T | TMD2 | 790 ± 71 | 2275 ± 162 | 2381 ± 161 | 15 ± 126 | 187 ± 3 | 686 ± 17 | 667 ± 91 | 1736 ± 13 |
| **R1162L** | 3485G>T | NBD2 | 828 ± 88 | 2243 ± 150 | 2338 ± 176 | 4138 ± 273 | 217 ± 7 | 754 ± 21 | 752 ± 82 | 1772 ± 4 |
| **I1234V** | 3700A>G | NBD2 | 1331 ± 52 | 3332 ± 189 | 3353 ± 128 | 5680 ± 377 | 362 ± 19 | 1303 ± 21 | 1157 ± 140 | 2659 ± 26 |
| **S1235R** | 3705T>G | NBD2 | 1345 ± 59 | 3405 ± 192 | 3378 ± 119 | 5751 ± 320 | 361 ± 14 | 1246 ± 33 | 1106 ± 108 | 2593 ± 91 |
| **V1240G** | 3719T>G | NBD2 | 671 ± 157 | 1840 ± 350 | 1793 ± 387 | 3072 ± 524 | 164 ± 14 | 644 ± 59 | 583 ± 95 | 1356 ± 49 |

| Variant | Amino Acid Substitution | CFTR sub-domain | Parental Surface Immuno-staining Intensity (Apo) | Parental Surface Immuno-staining Intensity (VX-661) | Parental Surface Immuno-staining Intensity (VX-445) | Parental Surface Immuno-staining Intensity (VX-445 + VX-661) | CANX KO Surface Immuno-staining Intensity (Apo) | CANX KO Surface Immuno-staining Intensity (VX-661) | CANX KO Surface Immuno-staining Intensity (VX-445) | CANX KO Surface Immuno-staining Intensity (VX-445 + VX-661) |
|---|---|---|---|---|---|---|---|---|---|---|
| **G1244E** | 3731G>A | NBD2 | 1352 ± 23 | 3364 ± 206 | 3353 ± 184 | 5631 ± 384 | 375 ± 21 | 1290 ± 11 | 1176 ± 130 | 2727 ± 97 |
| **T1246I** | 3737C>T | NBD2 | 1293 ± 59 | 3253 ± 144 | 3286 ± 87 | 5633 ± 307 | 335 ± 11 | 1133 ± 45 | 1044 ± 109 | 2439 ± 72 |
| **G1249R** | 3745G>A | NBD2 | 1220 ± 51 | 3155 ± 176 | 3087 ± 75 | 5352 ± 348 | 357 ± 20 | 1248 ± 38 | 1153 ± 154 | 2727 ± 141 |
| **S1251N** | 3752G>A | NBD2 | 1353 ± 41 | 3290 ± 176 | 3313 ± 165 | 5769 ± 330 | 371 ± 17 | 1276 ± 8 | 1146 ± 129 | 2654 ± 26 |
| **S1255P** | 3763T>C | NBD2 | 1258 ± 54 | 3153 ± 173 | 3161 ± 89 | 5435 ± 299 | 318 ± 15 | 1136 ± 36 | 1025 ± 121 | 2374 ± 16 |
| **I1269N** | 3806T>A | NBD2 | 554 ± 106 | 1420 ± 324 | 1419 ± 344 | 2391 ± 432 | 149 ± 4 | 547 ± 16 | 492 ± 47 | 1169 ± 7 |
| **D1270N** | 3808G>A | NBD2 | 1242 ± 88 | 3122 ± 51 | 3151 ± 42 | 5479 ± 180 | 342 ± 12 | 1199 ± 29 | 1093 ± 122 | 2573 ± 44 |
| **W1282R** | 3844T>C | NBD2 | 363 ± 35 | 925 ± 264 | 949 ± 230 | 1551 ± 393 | 110 ± 16 | 326 ± 32 | 302 ± 43 | 720 ± 22 |
| **R1283M** | 3848G>T | NBD2 | 422 ± 59 | 1132 ± 308 | 1151 ± 270 | 1855 ± 418 | 122 ± 16 | 398 ± 38 | 375 ± 42 | 843 ± 15 |
| **R1283S** | 3849G>C | NBD2 | 465 ± 82 | 1252 ± 336 | 1220 ± 290 | 2064 ± 431 | 130 ± 15 | 430 ± 15 | 414 ± 52 | 982 ± 43 |
| **Q1291R** | 3872A>G | NBD2 | 1389 ± 70 | 3474 ± 173 | 3453 ± 110 | 5869 ± 326 | 361 ± 16 | 1275 ± 15 | 1120 ± 131 | 2516 ± 78 |
| **Q1291H** | 3873G>C | NBD2 | 1280 ± 62 | 3203 ± 104 | 3235 ± 86 | 5568 ± 203 | 327 ± 14 | 1083 ± 25 | 1018 ± 111 | 2285 ± 55 |
| **V1293G** | 3878T>G | NBD2 | 1331 ± 61 | 3329 ± 164 | 3314 ± 47 | 5581 ± 117 | 363 ± 14 | 1250 ± 23 | 1119 ± 128 | 2487 ± 47 |
| **N1303K** | 3909C>G | NBD2 | 262 ± 34 | 630 ± 154 | 609 ± 58 | 1034 ± 219 | 88 ± 15 | 226 ± 6 | 206 ± 30 | 432 ± 5 |
| **L1324P** | 3971T>C | NBD2 | 249 ± 44 | 579 ± 124 | 548 ± 31 | 975 ± 211 | 91 ± 14 | 223 ± 7 | 210 ± 16 | 441 ± 18 |
| **L1335P** | 4004T>C | NBD2 | 243 ± 30 | 623 ± 151 | 564 ± 57 | 967 ± 168 | 97 ± 15 | 224 ± 25 | 224 ± 13 | 458 ± 5 |
| **G1349D** | 4046G>A | NBD2 | 1225 ± 38 | 3029 ± 77 | 3062 ± 59 | 5304 ± 214 | 338 ± 20 | 1152 ± 46 | 1060 ± 115 | 2437 ± 57 |
| **I1366N** | 4097T>A | NBD2 | 408 ± 59 | 1050 ± 265 | 1048 ± 239 | 1685 ± 374 | 148 ± 11 | 558 ± 22 | 510 ± 79 | 1163 ± 50 |
| **H1375P** | 4124A>C | | 1253 ± 69 | 3167 ± 42 | 3171 ± 66 | 5497 ± 218 | 353 ± 31 | 1287 ± 10 | 1171 ± 126 | 2767 ± 134 |
| **L1480P** | 4439T>C | | 1080 ± 93 | 2698 ± 85 | 2658 ± 60 | 4535 ± 131 | 295 ± 12 | 1098 ± 38 | 941 ± 123 | 2181 ± 84 |
| **L138ins** | 413_415dupTAC | TMD1 | 201 ± 49 | 572 ± 115 | 486 ± 40 | 2039 ± 293 | 75 ± 14 | 171 ± 20 | 142 ± 18 | 579 ± 25 |
| **F312DEL** | 935_937del | NBD1 | 647 ± 77 | 2008 ± 224 | 2329 ± 124 | 4508 ± 98 | 164 ± 2 | 683 ± 24 | 762 ± 87 | 1940 ± 156 |
| **I507del** | 1519_1521delATC | NBD1 | 199 ± 51 | 425 ± 87 | 365 ± 50 | 626 ± 89 | 75 ± 18 | 123 ± 14 | 112 ± 21 | 211 ± 29 |
| **F508del** | 1521_1523delCTT | NBD1 | 195 ± 47 | 453 ± 108 | 410 ± 47 | 891 ± 150 | 68 ± 15 | 110 ± 12 | 110 ± 16 | 256 ± 26 |
| **T854T** | 2562T>A | | 1294 ± 51 | 3240 ± 145 | 3213 ± 88 | 5519 ± 358 | 357 ± 10 | 1216 ± 17 | 1072 ± 125 | 2486 ± 54 |
| | 2562T>C | | 1329 ± 72 | 3214 ± 194 | 3270 ± 21 | 5657 ± 238 | 350 ± 18 | 1193 ± 54 | 1063 ± 117 | 2478 ± 68 |
| | 2562T>G | | 1287 ± 53 | 3169 ± 189 | 3164 ± 147 | 5401 ± 312 | 364 ± 14 | 1251 ± 44 | 1134 ± 116 | 2588 ± 72 |
| **Q966=** | 2988G>A | TMD2 | 1324 ± 43 | 3171 ± 147 | 3196 ± 88 | 5527 ± 156 | 332 ± 19 | 1132 ± 59 | 1000 ± 95 | 2308 ± 68 |
| **L1156=** | 3598T>C-3466T>C | TMD2 | 1299 ± 58 | 3298 ± 269 | 3167 ± 139 | 5429 ± 258 | 373 ± 24 | 1255 ± 5 | 1117 ± 96 | 2651 ± 138 |
| | 3600G>A-3468G>A | TMD2 | 1308 ± 55 | 3307 ± 201 | 3213 ± 160 | 5414 ± 261 | 373 ± 25 | 1299 ± 31 | 1151 ± 136 | 2695 ± 105 |
| | 3468G>A | TMD2 | 1312 ± 46 | 3232 ± 178 | 3186 ± 149 | 5514 ± 255 | 346 ± 16 | 1162 ± 21 | 1037 ± 77 | 2386 ± 60 |
| **E60X** | 178G>T | Lasso | 207 ± 46 | 478 ± 136 | 393 ± 33 | 675 ± 93 | 79 ± 14 | 148 ± 29 | 130 ± 23 | 272 ± 48 |
| **Q493X** | 1477C>T | NBD1 | 198 ± 43 | 412 ± 73 | 384 ± 56 | 647 ± 97 | 69 ± 16 | 113 ± 17 | 107 ± 20 | 196 ± 34 |
| **G542X** | 1624G>T | NBD1 | 205 ± 40 | 459 ± 104 | 403 ± 43 | 652 ± 76 | 73 ± 19 | 117 ± 19 | 109 ± 15 | 205 ± 46 |
| **R553X** | 1657C>T | NBD1 | 203 ± 38 | 442 ± 77 | 402 ± 17 | 647 ± 80 | 69 ± 14 | 116 ± 21 | 106 ± 18 | 199 ± 33 |
| **S912X** | 2735C>A | TMD2 | 202 ± 49 | 428 ± 82 | 387 ± 35 | 664 ± 128 | 74 ± 16 | 125 ± 17 | 115 ± 21 | 225 ± 43 |
| **W1089X** | 3266G>A | TMD2 | 206 ± 46 | 448 ± 105 | 396 ± 51 | 690 ± 96 | 71 ± 16 | 124 ± 13 | 110 ± 14 | 198 ± 34 |
| **Y1092X** | 3276C>A | TMD2 | 210 ± 39 | 457 ± 112 | 402 ± 44 | 663 ± 67 | 70 ± 14 | 134 ± 14 | 118 ± 21 | 227 ± 22 |
| | 3276C>G | TMD2 | 210 ± 49 | 490 ± 132 | 418 ± 43 | 689 ± 124 | 74 ± 14 | 129 ± 21 | 124 ± 25 | 237 ± 37 |
| **E1104X** | 3310G>T | TMD2 | 195 ± 49 | 453 ± 159 | 390 ± 50 | 675 ± 118 | 68 ± 15 | 114 ± 12 | 104 ± 19 | 192 ± 31 |
| **R1158X** | 3472C>T | TMD2 | 220 ± 56 | 523 ± 135 | 447 ± 81 | 740 ± 110 | 86 ± 16 | 189 ± 11 | 152 ± 13 | 351 ± 26 |
| **R1162X** | 3484C>T | TMD2 | 206 ± 39 | 463 ± 108 | 402 ± 39 | 696 ± 90 | 73 ± 15 | 130 ± 15 | 129 ± 29 | 256 ± 10 |
| **S1196X** | 3587C>G | TMD2 | 962 ± 166 | 2220 ± 202 | 2055 ± 254 | 3143 ± 290 | 245 ± 8 | 856 ± 26 | 701 ± 95 | 1381 ± 57 |
| **W1204X** | 3611G>A | NBD2 | 971 ± 147 | 2170 ± 251 | 1897 ± 278 | 2866 ± 417 | 234 ± 5 | 806 ± 54 | 622 ± 75 | 1265 ± 32 |
| | 3612G>A | NBD2 | 916 ± 167 | 2002 ± 339 | 1736 ± 319 | 2472 ± 407 | 214 ± 9 | 690 ± 80 | 543 ± 75 | 1045 ± 33 |

| Variant | Amino Acid Substitution | CFTR sub-domain | Parental Surface Immuno-staining Intensity (Apo) | Parental Surface Immuno-staining Intensity (VX-661) | Parental Surface Immuno-staining Intensity (VX-445) | Parental Surface Immuno-staining Intensity (VX-445 + VX-661) | CANX KO Surface Immuno-staining Intensity (Apo) | CANX KO Surface Immuno-staining Intensity (VX-661) | CANX KO Surface Immuno-staining Intensity (VX-445) | CANX KO Surface Immuno-staining Intensity (VX-445 + VX-661) |
|---|---|---|---|---|---|---|---|---|---|---|
| **W1282X** | 3846G>A | NBD2 | 353 ± 29 | 908 ± 247 | 900 ± 149 | 1533 ± 295 | 108 ± 8 | 365 ± 3 | 332 ± 64 | 825 ± 146 |
| **Q1313X** | 3937C>T | NBD2 | 202 ± 52 | 461 ± 126 | 404 ± 50 | 668 ± 108 | 68 ± 16 | 113 ± 21 | 106 ± 22 | 187 ± 32 |
